# Decoding the genomic landscape of chromatin-associated biomolecular condensates

**DOI:** 10.1101/2023.08.23.554542

**Authors:** Zhaowei Yu, Qi Wang, Guangya Zhu, Jidong Zhu, Yong Zhang

## Abstract

Biomolecular condensates play a significant role in chromatin activities, primarily by concentrating and compartmentalizing proteins and/or nucleic acids. However, their genomic landscapes and compositions remain largely unexplored due to a lack of dedicated computational tools for systematic identification *in vivo*. To address this, we developed CondSigDetector, a computational framework designed to detect condensate-like chromatin-associated protein co-occupancy signatures (CondSigs), to predict genomic loci and component proteins of distinct chromatin-associated biomolecular condensates. Applying this framework to mouse embryonic stem cells (mESC) and human K562 cells enabled us to depict the high-resolution genomic landscape of chromatin-associated biomolecular condensates, and uncover both known and potentially novel biomolecular condensates. Multi-omics analysis and experimental validation further verified the condensation properties of CondSigs. Additionally, our investigation shed light on the impact of chromatin-associated biomolecular condensates on chromatin activities. Collectively, CondSigDetector provides a novel approach to decode the genomic landscape of chromatin-associated condensates, facilitating a deeper understanding of their biological functions and underlying mechanisms in cells.

## Introduction

Over the last decade, there has been growing appreciation for the biological role of biomolecular condensates, which are membraneless compartments that compartmentalize and concentrate specific proteins and/or nucleic acids^1,2^. Liquid-liquid phase separation (LLPS) has been proposed as a key organizing principle of biomolecular condensates, driven by weak, multivalent, and highly collaborative molecular interactions^2^. The molecular interactions inside biomolecular condensates usually involve diverse collaborative components that can be categorized into two main groups: scaffolds and clients. Scaffolds drive the formation of condensates, while clients participate by binding to scaffolds^3–6^. Biomolecular condensates are implicated in various cellular functions, and their aberrations are associated with numerous diseases^1,7^. Recently, growing evidences have demonstrated the widespread existence and functional significance of chromatin-associated biomolecular condensates. Many chromatin-associated processes, such as DNA replication^8^, DNA repair^9^, transcription control^10–13^, and chromatin organization^14–17^, have been found to take place within biomolecular condensates at chromatin^18^ (Supplementary Table S1).

Understanding chromatin-associated biomolecular condensates, including their genomic loci and collaborative components, is crucial for elucidating their impact on chromatin activities. Although some chromatin-associated biomolecular condensates have been linked to well-characterized chromatin states, such as super-enhancer^10,11^ and heterochromatin^15–17^, these connections have generally been reported without comprehensive associations with genome-wide loci, except for a few loci of interest validated by low-throughput experiments. Until now, the genomic landscape of chromatin-associated biomolecular condensates has remained poorly understood. However, no genomic approach has been designed yet to capture the comprehensive genomic landscape of chromatin-associated biomolecular condensates, primarily due to the following challenges. First, the complexity of biomolecular condensates arising from their diverse components^18^ and context-specific molecular collaborations among these components along the chromatin^13^, making it difficult to systematically capture chromatin-associated biomolecular condensates by targeting a single factor. Second, even for chromatin-associated protein (CAP) with experimental evidence of condensation^3–6,19^, distinguishing its condensation-associated binding sites from non-associated binding sites in individual datasets is not a straightforward task.

With the rapid accumulation of CAP occupancy profiles and proteome-scale characterization of condensation potential, it is now possible to overcome the above challenges of decoding the genomic landscape of chromatin-associated biomolecular condensates by integrating multi-dimensional data. In this study, we introduce CondSigDetector, a computational framework that systematically predicts chromatin-associated biomolecular condensates. This framework overcomes the two challenges mentioned above by utilizing topic modeling to detect genome-wide context-dependent collaborations among CAPs possessing high condensation potential from hundreds of CAP occupancy profiles. These collaborations along the chromatin are termed <u>Cond</u>ensate-like chromatin-associated protein co-occupancy <u>Sig</u>natures (CondSigs). The framework not only identifies the collaborative components of distinct biomolecular condensates, but also assigns them to the associated genomic loci. We applied this computational framework to two cell types with abundant ChIP-seq data, and predicted hundreds of chromatin-associated biomolecular condensates, along with their genomic loci, which are supported by multi-omics data and experimental evidences. To the best of our knowledge, CondSigDetector is the first computational framework for decoding the genomic landscape of chromatin-associated biomolecular condensates, providing a valuable resource for investigating the functional effects and underlying mechanisms of chromatin-associated biomolecular condensates on chromatin activities.

## Results

### Overall design of CondSigDetector

By integrating ChIP-seq datasets of hundreds of CAPs in the same cell type, we observed frequent co-occupancy of CAPs across the genome (Supplementary Fig. S1a, b). However, most co-occupancy events could not be explained by DNA binding motifs or chromatin accessibility (Supplementary Fig. S1c-f), two known determinants of CAP co-occupancy events^20^. This suggests that alternative mechanisms may be responsible for organizing genome-wide co-occupancy events of CAPs. Biomolecular condensation at chromatin may partially explain such events, as biomolecular condensates are thought to be mediated by collaborations of components^2^, and condensations of CAPs have been reported to influence their chromatin occupancy^10,21^. This evidence implies that specific CAP co-occupancy events could be signatures of chromatin-associated biomolecular condensates.

In this study, we aim to predict chromatin-associated biomolecular condensates by detecting genome-wide context-dependent collaborations of CAPs with high condensation potential, termed CondSig. We developed a computational framework, CondSigDetector, to systematically detect CondSigs by integrating hundreds of ChIP-seq datasets and condensation-related characterizations of CAPs (Fig. 1). CondSigDetector comprises three steps: data processing, co-occupancy signatures identification, and condensation potential filtration.

**Fig. 1.**
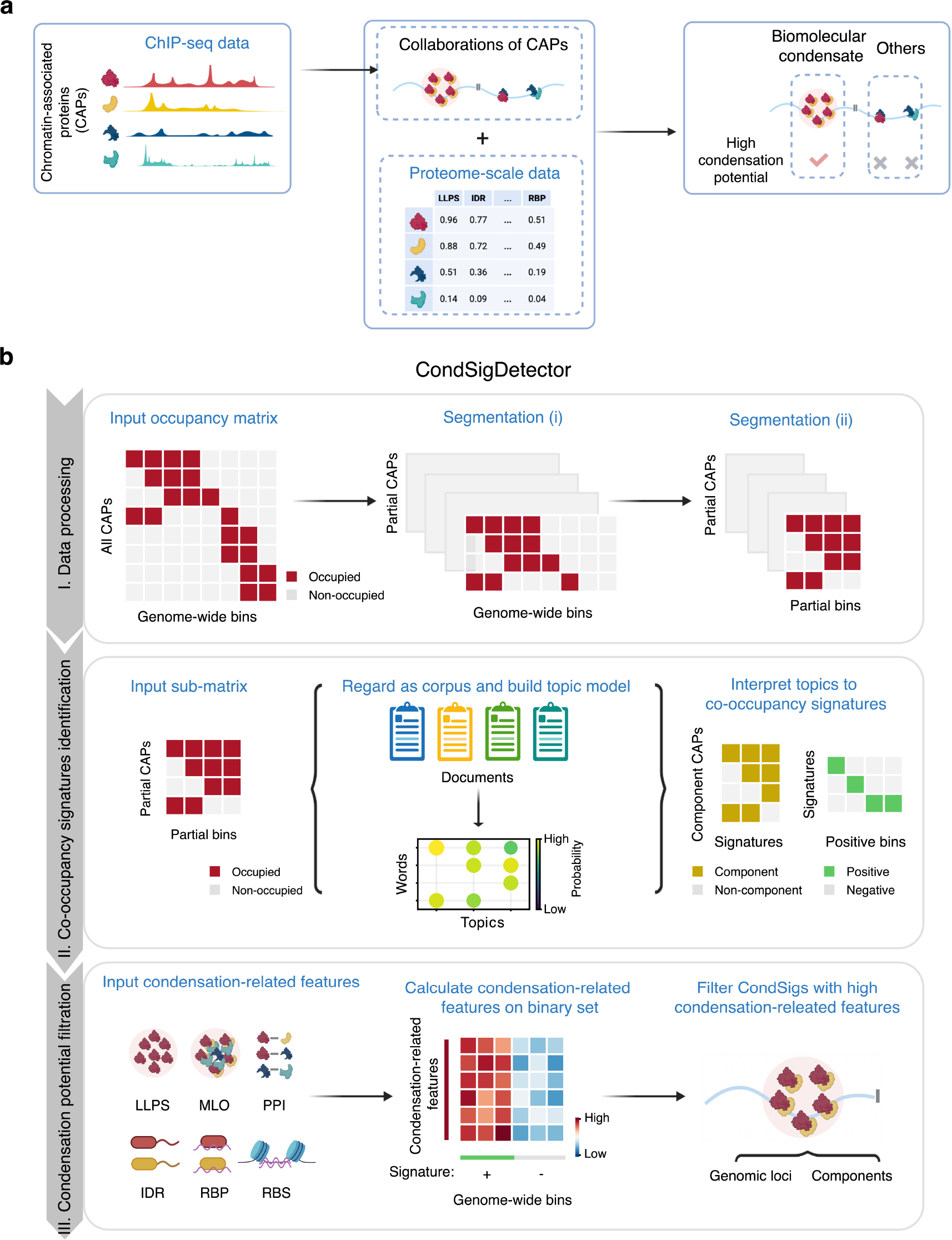
Overall design of CondSigDetector. **a.** A schematic illustrates the prediction of chromatin-associated biomolecular condensates by detecting genome-wide context-dependent collaborations of CAPs with high condensation potential. **b.** Workflow of CondSigDetector (see Methods for details).

In the first step, the input data, *i.e.*, the collected ChIP-seq profiles of all CAPs from an identical cell type, is converted into an occupancy matrix at genome-wide consecutive bins. To address the sparsity of this matrix, CondSigDetector applies an iterative segmentation method for each target CAP, which segments the entire occupancy matrix into smaller sub-matrices (see Methods for details). This segmentation approach can enhance the detection of CAP collaborations in local contexts by substantially increasing the occurrence frequency of co-occupancy events within the sub-matrices (Supplementary Fig. S1g, h).

In the second step, CondSigDetector utilizes a topic model to identify co-occupancy signatures of CAPs, representing frequent CAP collaborations, from the sub-matrices. Given the significant differences in co-occupancy frequencies between promoter and non-promoter regions (Supplementary Fig. S1a, b), the sub-matrices are categorized into either promoter or non-promoter groups to identify co-occupancy signatures separately. Within the topic model, each sub-matrix is treated as a set of documents, where each genomic bin represents a document and CAPs occupying the bin are considered as words in the document. Intuitively, the topics learned from topic modeling, which indicate specific word combinations, can be interpreted as co-occupancy signatures of CAPs. Since the number of co-occupied CAPs within a bin is typically sparse (Supplementary Fig. S1a, b), CondSigDetector utilizes the biterm topic model, which outperforms traditional models such as Latent Dirichlet Allocation for short text^22^. It has been confirmed that the co-occupancy signatures of CAPs derived from the biterm topic model exhibit high topic coherence and repeatability among replicates (see Methods for details; Supplementary Fig. S1i-l).

In the third step, CondSigDetector predicts CondSigs by evaluating the condensation potential for each co-occupancy signature of CAPs. For each genomic bin, 6 condensation-related features are calculated: the fraction of occupied CAPs with reported LLPS capacity, the fraction of occupied CAPs co-occurring in the same membraneless organelle (MLO), the fraction of occupied CAPs with predicted intrinsically disordered regions (IDRs), the fraction of occupied CAP pairs having protein-protein interactions (PPIs), the fraction of occupied CAPs predicted as RNA-binding proteins (RBPs), and the RNA-binding strength (RBS) of the bin. Intuitively, for a co-occupancy signature of CAPs, higher values of these condensation-related features at signature-positive bins indicate a greater condensation potential. Co-occupancy signatures with at least 3 condensation-related features strongly and positively correlated with their presence are identified as CondSigs (see Methods for details). Finally, CondSigDetector eliminates redundant CondSigs containing similar CAP components.

### Identification of CondSigs in mouse and human cell lines

CondSigDetector was applied to two cell types with abundant ChIP-seq data: mESC and human K562 cell line, to identify CondSigs. After stringent quality control, we gathered qualified ChIP-seq data for 189 CAPs in mESC and 216 CAPs in K562 (Supplementary Table S2). Due to the lack of a qualified RNA-binding profile for mESC, the RNA binding strength, one of the condensation-related features, was not included in mESC. We identified 25 promoter CondSigs and 36 non-promoter CondSigs in mESC (Fig. 2a), along with 75 promoter CondSigs and 93 non-promoter CondSigs in K562 (Supplementary Fig. S2). Additionally, we identified 14,345 promoter CondSig-positive sites and 24,500 non-promoter CondSig-positive sites in mESC, along with 14,201 and 38,963 CondSig-positive sites in K562. To assess the reliability of identified CondSigs, we examined whether their component CAPs are involved in known chromatin-associated biomolecular condensates. Among the identified mESC CondSigs, 92.0% of promoter and 97.2% of non-promoter CondSigs contain at least one component CAP present in known chromatin-associated biomolecular condensates (Fig. 2b). For example, a non-promoter CondSig contains SS18, SMARCA4 (BRG1), and DPF2, which are three known components of the known SS18 cluster^23^ (Fig. 2a, Supplementary Fig. S3a, Supplementary Table S1). In K562 cells, 49.3% of promoter and 55.9% of non-promoter CondSigs have at least one component CAP found in known chromatin-associated biomolecular condensates (Fig. 2b). One example of a non-promoter CondSig includes CBX5 (HP1α), TRIM28 and CBX1 (HP1β) (Supplementary Fig. S3b, Supplementary Table S1), with HP1 and TRIM28 were reported to drive LLPS with H3K9me3-modified chromatin cooperatively^15^. These results provide support for the reliability of the identified CondSigs.

**Fig. 2.**
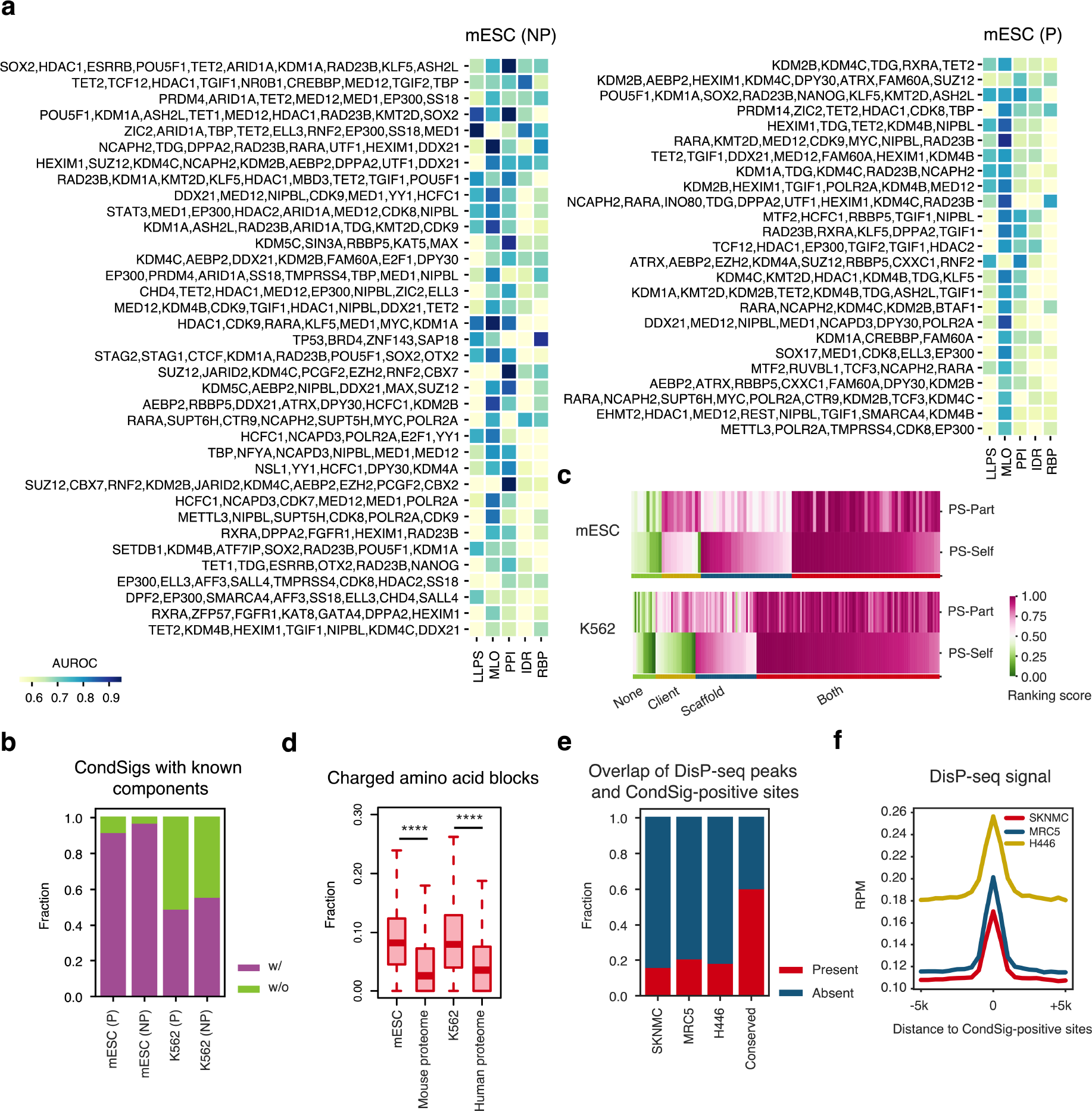
CondSigs in mESC and K562. **a.** Heatmaps showing identified CondSigs in mESC at non-promoter (NP) and promoter (P) regions. Each row represents a CondSig and the row name indicates the component CAPs of the given CondSig. Each column represents a condensation-related feature and the colours represent AUROC. **b.** The stacked bar plots showing the fraction of CondSig with/without components in known chromatin-associated biomolecular condensates (Supplementary Table S1). **c.** Heatmaps showing *k*-means clustering for component CAPs in mESC (**left**) and K562 (**right**) with PS-self and PS-part ranking score (see Methods for details). A high PS-self ranking score refers to scaffolds and a high PS-part ranking score refers to clients. Four clusters (“Both”: both scaffold and client, “Scaffold”: scaffold-only, “Client”: client-only, and “None”) were shown. **d.** Box plots showing component CAPs of CondSigs have a higher fraction of charged blocks in amino acid sequences (see Methods for details) than control (mouse or human proteome) in mESC and K562. Significance between groups was evaluated by a two-sided Welch’s *t*-test, **** represents *p*-value < 1 × 10^−4^. **e.** The overlap ratio of DisP-seq peaks from three human cell lines with CondSig-positive sites in K562. Conserved peaks represent shared peaks across three human cell lines. DisP-seq data were from the previous study (GSE190961^28^). **f.** Line chart showing average DisP-seq signals around CondSig-positive sites. *X*-axis represents distance to CondSig-positive site centers and *Y*-axis represents average DisP-seq signals calculated by RPM (reads per million mapped reads).

Some component CAPs are found in more than one identified CondSig (Supplementary Fig. S3c-d). For example, DDX21, a DEAD-box RNA helicase known to participate in biomolecular condensate^24^, is present in 8 non-promoter CondSigs in mESC. We examined the similarity of present loci between CondSig pairs containing at least one shared component CAP and found that only 0.6% of mESC pairs and 0.9% of K562 pairs had a Jaccard index higher than 0.7. This suggests a high diversity of present loci of identified CondSigs, even when they share some common components. To investigate the potential roles of component CAPs in CondSigs, we classified all predicted component CAPs into four clusters: “both scaffold and client”, “scaffold-only”, “client-only”, and “none”, according to their calculated potentials for self-assembly or interaction with partners to undergo phase separation^25^ (see Methods for details). 77.5% and 79.5% of component CAPs in mESC and K562 were classified into “both scaffold and client” or “scaffold-only” clusters (Fig. 2c). Furthermore, we found that component CAPs of CondSigs have a significantly higher fraction of charged amino acid blocks (Fig. 2d), which is an important resource for multivalency^26^. These results demonstrate that the component CAPs of identified CondSigs have strong capacities to form biomolecular condensates, and may function in a context-dependent manner.

The previous studies demonstrated that biomolecular condensate can form at super-enhancers, *i.e.*, clusters of enhancers densely occupied by the master regulators and mediators, and these condensates can regulate gene transcription by concentrating transcription machinery^10,27^. When comparing the genomic loci of super-enhancers and CondSig-positive sites in mESC, we found that 96.1% of super-enhancers overlap with CondSig-positive sites. Furthermore, a recent study introduced DisP-seq, an antibody-independent chemical precipitation assay, to map genome-wide profiles of disordered proteins^28^. We reanalyzed public DisP-seq data from three human cell lines and compared the DisP-seq peaks with identified CondSig-positive sites in the K562 cell line. In SKNMC, MRC5, and H446 cells, 16.1%, 21.0%, and 18.5% of DisP-seq peaks, respectively, were identified as CondSig-positive sites in K562. But among the shared DisP-seq peaks across the three human cell lines, 60.6% were identified as CondSig-positive sites in K562 (Fig. 2e). We further observed much higher DisP-seq signals at CondSig-positive sites in K562 compared to their adjacent regions (Fig. 2f), suggesting that identified CondSig-positive sites are highly occupied by disordered proteins, which have been demonstrated to play important roles in biomolecular condensation^2^. These results point towards the high potential of identified CondSig-positive sites as genomic loci where biomolecular condensates form.

### Chromatin properties of identified CondSigs

To investigate the chromatin features of identified CondSigs, we first analyzed the concentration levels of the component within CondSigs by calculating ChIP-seq signal strength for each component. We divided the ChIP-seq peaks of each component CAP into CondSig-positive groups and -negative groups based on their overlap with positive sites of CondSigs (see Methods for details), and compared their ChIP-seq signals. As shown in Fig. 3a, most component CAPs displayed significantly higher signal strength at CondSig-positive peaks in mESC, indicating that CondSigs can concentrate their components at target genomic loci. For example, CTCF, a CAP involved in chromatin insulation^29^, exhibited significantly higher signal strength at CondSig-positive CTCF peaks. To investigate the biological functional effect of CTCF concentration, we re-analyzed Micro-C data in mESC^30^ and found that CondSig-positive CTCF peaks exhibited significantly higher boundary strength than CondSig-negative CTCF peaks (Fig. 3b), suggesting that CTCF concentration contributes to enhanced chromatin insulation activity. We then merged the adjacent ChIP-seq peaks to obtain domains for each component CAP (see Methods for details), and compared the width distributions of CondSig-positive and -negative domains. As shown in Fig. 3c, CondSig-positive domains are wider on average for 95.2% and 93.5% of all component CAPs of promoter and non-promoter CondSigs, and the CondSig-positive domains of RUVBL1, TCF3, CTR9, MTF2 and SUPT6H exceeded 10 kb on average. Additionally, we assessed the component concentration levels and domain widths of CondSigs in K562 and found largely consistent results (Supplementary Fig. S4a, b). These results confirmed the component concentration properties of CondSigs, which is a basic feature of known chromatin-associated biomolecular condensates^18^, and suggested a potential association between biomolecular condensation and stronger effects on chromatin activities.

**Fig. 3.**
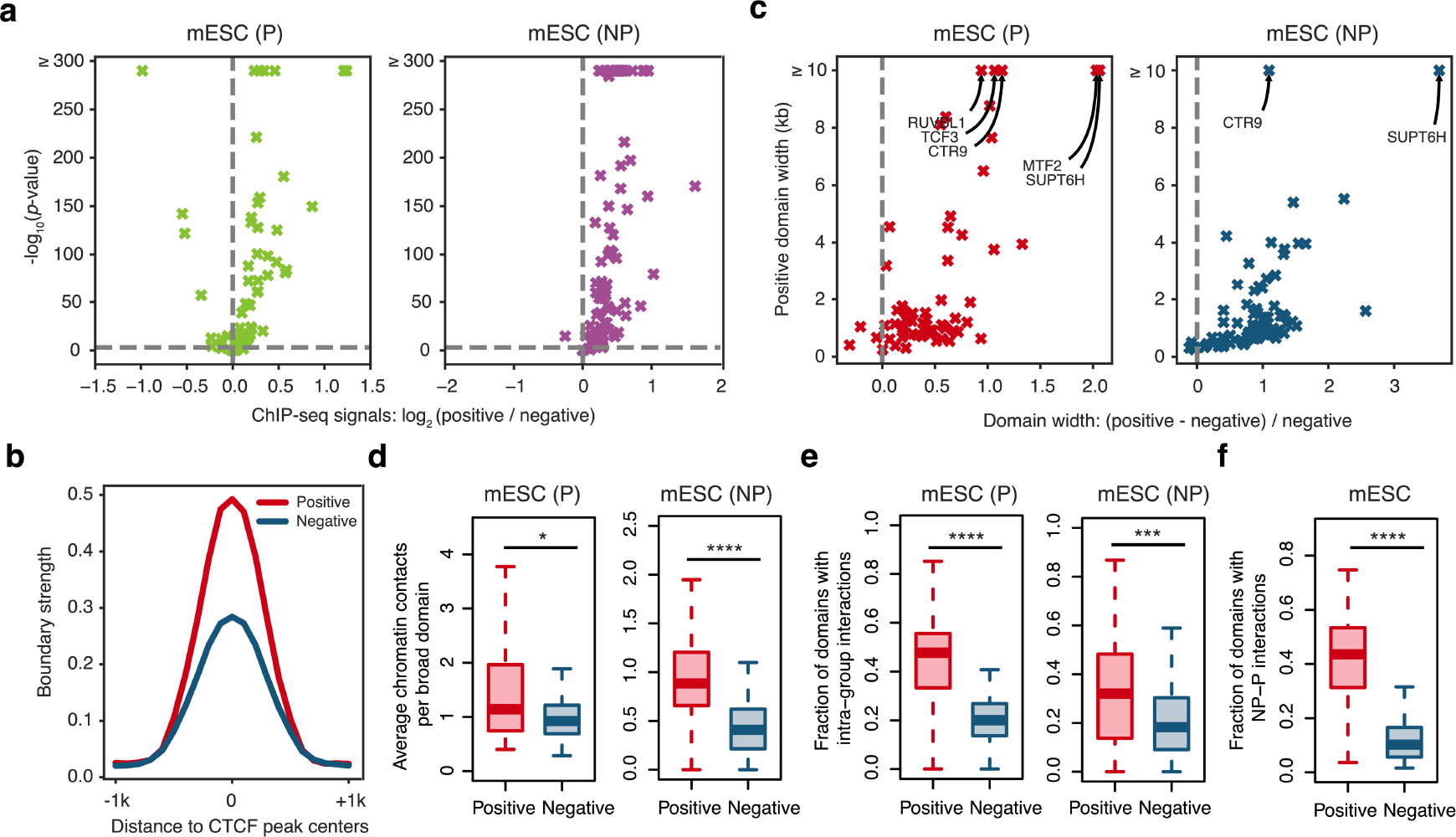
Chromatin properties of predicted chromatin-associated biomolecular condensates. **a.** Volcano plots showing concentration levels of component CAPs in CondSigs in mESC. *X*-axis represents the log_2_-transformed fold change of ChIP-seq signals at CondSig-positive peaks compared to CondSig-negative peaks, while *Y*-axis represents the negative log_10_-transformed *p*-value. The vertical dashed line corresponds fold change = 1 and the horizontal dashed line corresponds to *p*-value = 0.001. **b.** Line charts showing boundary strength around CondSig-positive and - negative CTCF peaks. Boundary strength was calculated using Micro-C data in mESC from the previous study (GSE130275^30^). **c.** Scatter plots showing width comparison of CondSig-positive and -negative domains in mESC. *X*-axis represents the ratio to which the CondSig-positive domain width exceeds the CondSig-negative domain width, and *Y*-axis represents the positive domain width. Component CAPs having CondSig-positive domains exceeding 10 kb on average were labeled. **d.** Intra-domain chromatin contacts of CondSig-positive or -negative broad domains in mESC. For each component CAP, an average valid paired-end tags count in each broad domain (> 5 kb) was calculated to represent intra-domain contacts. Cohesin ChIA-PET data used in the analysis was from the previous study (GSE57913^33^). **e.** Box plots showing intra-group chromatin contacts between CondSig-positive or - negative domains in mESC. For each component CAP, the fraction of domains having at least one valid paired-end tag with other intra-group domains was calculated. **f.** Box plots showing NP (non-promoter)-P (promoter) chromatin contacts between CondSig-positive or -negative domains in mESC. For each component CAP, the fraction of non-promoter domains having at least one valid paired-end tag with its promoter domains was calculated. Significance between groups was evaluated by a two-sided Welch’s *t*-test, * represents *p*-value < 0.05, *** represents *p*-value < 1 × 10^−3^ and **** represents *p*-value < 1 × 10^−4^.

Based on previous studies that reported spatially proximal chromatin could be involved in the same condensates^31,32^, we processed to analyze chromatin contact frequencies within and between CondSig-positive and -negative domains for each component CAP. In order to minimize the impact of distinct width distributions between CondSig-positive and -negative domains, we focused on broad domains (width > 5 kb). We used cohesin ChIA-PET data from mESC^33^ to measure chromatin interactions between genomic loci, and found that CondSig-positive domains exhibited significantly higher intra-domain interactions than their CondSig-negative counterparts (Fig. 3d). We further calculated the fractions of domains with chromatin interactions within the same group of domains for each component CAP, and found significantly higher frequencies between CondSig-positive domains compared to CondSig-negative domains (Fig. 3e). For each component CAP presented in both promoter and non-promoter CondSigs, we calculated fractions of domains with chromatin interactions between its promoter and non-promoter domains. Our analysis observed that CondSig-positive domains showed significantly higher frequencies between promoter and non-promoter domains relative to CondSig-negative domains (Fig. 3f). We also utilized Pol II ChIA-PET data^34^ to evaluate the chromatin contact frequencies of CondSigs in K562, and observed largely consistent results (Supplementary Fig. S4c-e). These results confirmed that the components of identified CondSigs can be concentrated in trans through spatially proximal chromatin.

### Involvement of DDX21 in chromatin-associated biomolecular condensate

Although DDX21 can undergo phase separation and has been reported to participate in nucleolar condensate for Pol I transcription^24,35^, additional genomic loci where it may involve into biomolecular condensate remain to be elucidated. In mESC, we identified 10 CondSigs with DDX21 as a component, with 15,578 DDX21 ChIP-seq peaks as CondSig-positive. To verify the presence of DDX21-associated biomolecular condensate at these genomic loci, we assessed the sensitivity of DDX21 occupancy at these loci to 1,6-hexanediol (1,6-HD), a compound used for disrupting liquid-like biomolecular condensates^36^. Cleavage Under Targets and Release Using Nuclease (CUT&RUN) experiments were conducted for DDX21 in both wild type and 1,6-HD-treated mESC. We observed a significantly greater decrease in DDX21 CUT&RUN signals at CondSig-positive peaks compared to CondSig-negative peaks (Fig. 4a, b), which demonstrated the strong effect of biomolecular condensate disruption on CondSig-positive peaks of DDX21. This result supported that DDX21 participates in biomolecular condensates at these loci. We further investigated the potential impact of DDX21-associated biomolecular condensates at these genomic loci. We found that target genes of the CondSig-positive peaks of DDX21 displayed significantly higher expression levels than other genes (Supplementary Fig. S5a), suggesting that DDX21-associated biomolecular condensate may enhance the transcription of target genes.

**Fig. 4.**
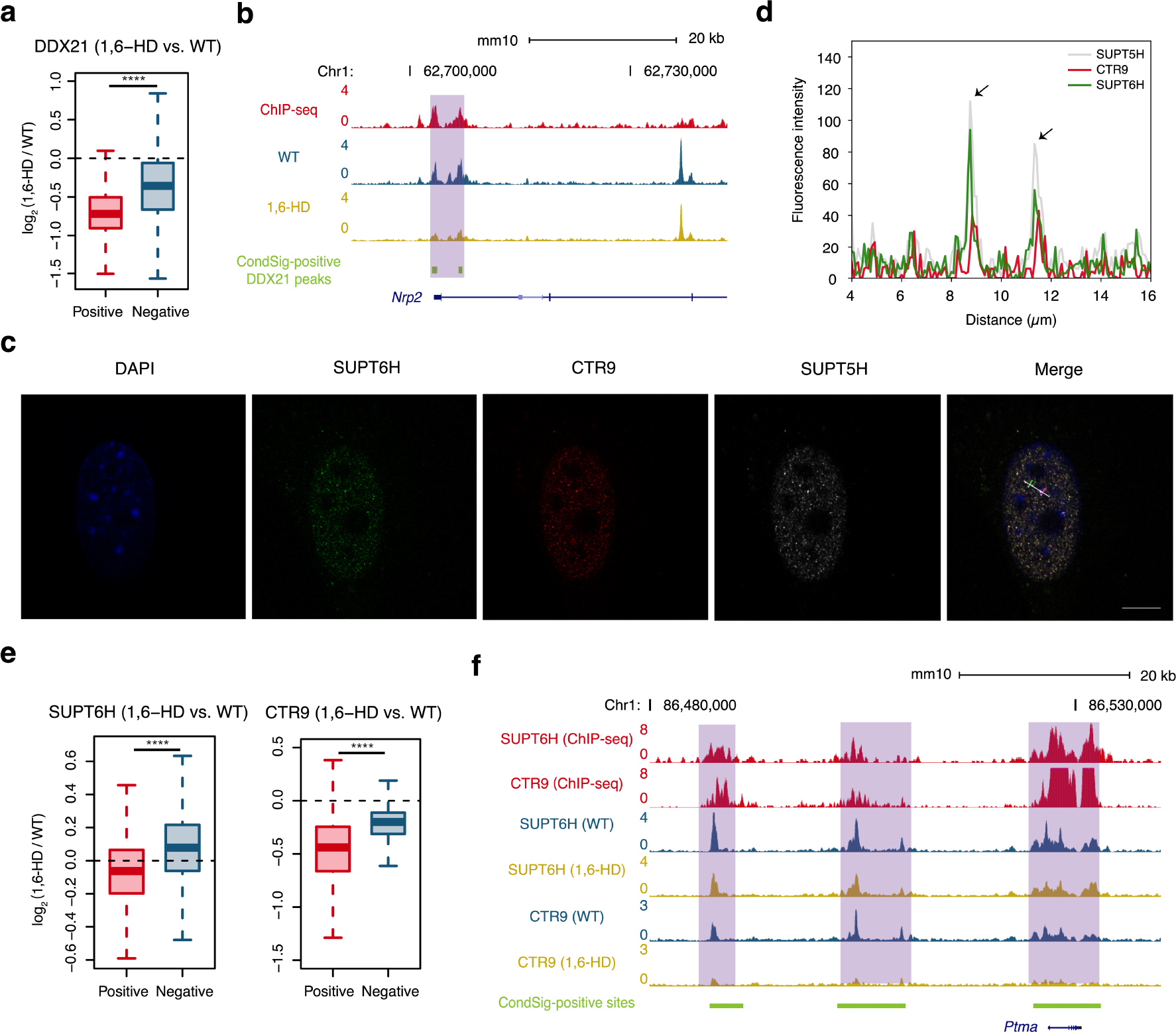
Experimental validation of potential biomolecular condensates. **a.** Box plots showing log_2_-transformed fold change of DDX21 CUT&RUN signals at CondSig-positive and -negative DDX21 peaks after 1,6-hexanediol in contrast to wild type mESC. **b.** UCSC genome browser view of representative CondSig-positive DDX21 peaks. Signals represent RPM and the related CondSig-positive site was shaded in purple. **c.** Immunofluorescence images of mESC showing that SUPT6H (green) colocalizes with CTR9 (red) and SUPT5H (grey) in puncta. DNA was stained with DAPI (blue). Scale bar: 10 *μ*m. **d.** Line scans of the images of a cell co-stained for SUPT6H, CTR9 and SUPT5H, at the position depicted by the white line. The direction is from the green tick to the purple tick, and the two arrows refer to two representative puncta. **e.** Box plots showing log_2_-transformed fold change of SUPT6H and CTR9 CUT&RUN signals at CondSig-positive sites and -negative sites after 1,6-hexanediol treatment in contrast to wild type mESC. Significance between groups was evaluated by a two-sided Welch’s *t*-test, **** represents *p*-value < 1 × 10^−4^. **f.** UCSC genome browser view of representative CondSig-positive sites. Signals represent RPM and the related loci were shaded in purple.

### Confirmation of CondSigs regulating transcription elongation

SUPT6H, SUPT5H and CTR9 have been reported to regulate transcription elongation^37,38^, but it remains unclear whether these CAPs function in the form of condensate. In mESC, we identified two CondSigs containing all or at least three of SUPT6H, SUPT5H, CTR9, and POLR2A simultaneously (Fig. 2a, Supplementary Fig, S5b). Genomic enrichment analysis found that merged CondSig-positive sites of the two CondSigs were primarily located at promoters and gene bodies (especially at exons), and the associated gene bodies were enriched with H3K36me3 modification, a marker for actively transcribed genes (Supplementary Fig. S5c, d). This suggested that SUPT6H, SUPT5H and CTR9 might participate in the same biomolecular condensate to regulate transcription elongation. To confirm the condensation properties of these component CAPs, we performed fixed cell immunofluorescence (IF) with antibodies against SUPT6H, SUPT5H and CTR9 in mESC. We found that all three CAPs can form nuclear puncta in cells (Fig. 4c), which is consistent with a recent study showing the condensation properties of SUPT6H and CTR9 in cells^39^. To determine whether these CAPs coexist in the same puncta, we conducted co-IF analysis and found their high co-localization in nuclei (Fig. 4c, d). To further verify the presence of the associated biomolecular condensate at these CondSig-positive sites, we conducted CUT&RUN experiments for SUPT6H and CTR9 in both wild type and 1,6-HD-treated mESC. We observed that CondSig-positive sites exhibited significantly greater decreases in CUT&RUN signals for both SUPT6H and CTR9 compared to control sites upon 1,6-HD treatment (Fig. 4e, f). These results suggested that SUPT6H, SUPT5H and CTR9 can regulate transcription elongation by forming biomolecular condensate.

### Effects of biomolecular condensate on chromatin activities

With the availability of CondSig-positive sites, it is possible to investigate the influence of biomolecular condensates on chromatin activities at a genome-wide scale. Our initial analysis for histone modifications at CondSig-positive sites revealed a high enrichment of active histone modifications, such as H3K4me3 and H3K27ac, in both mESC and K562 (Fig. 5a), suggesting a close association between biomolecular condensates and chromatin activities. We predicted the target genes associated with the CondSig-positive sites (see Methods for details), discovering that these genes showed significantly higher expression levels in both mESC and K562 (Supplementary Fig. S6a, b). Given that transcriptional bursting is a common characteristic of gene expression^40^, and it was hypothesized that biomolecular condensation can influence the transcriptional bursting frequencies of target genes^41^, we generated single-cell RNA-seq data in wild type and 1,6-HD-treated mESC and K562, from which we inferred transcriptome-wide transcriptional bursting kinetics^42^. Among the genes with inferable transcriptional bursting kinetics, those associated with CondSig-positive sites exhibited significantly higher bursting frequencies in the wild type mESC and K562 (Fig. 5b, c). They also displayed a more substantial decrease in transcriptional bursting frequencies upon 1,6-HD treatment compared to other genes (Fig. 5d, e). After assigning genes associated with CondSig-positive sites to individual CondSig, we ranked the CondSigs in mESC according to the decrease level of transcriptional bursting frequencies upon 1,6-HD treatment. As shown in Fig. 5f, the CondSig containing PRDM4, ARID1A, TET2, MED12, MED1, EP300 and SS18 demonstrated the most substantial decrease, suggesting that these CAPs may form biomolecular condensation to enhance the transcriptional bursting frequencies of their target genes. On the contrary, the CondSig containing SUZ12, JARID2, KDM4C, PCGF2, EZH2, RNF2 and CBX7 had the most increase, consistent with their repressive roles in transcription regulation^43^. And target genes of CondSigs in K562 exhibited decreased burst frequency on average (Supplementary Fig. S6c). These results suggested that biomolecular condensation can regulate gene transcription by influencing transcriptional bursting frequency.

**Fig. 5.**
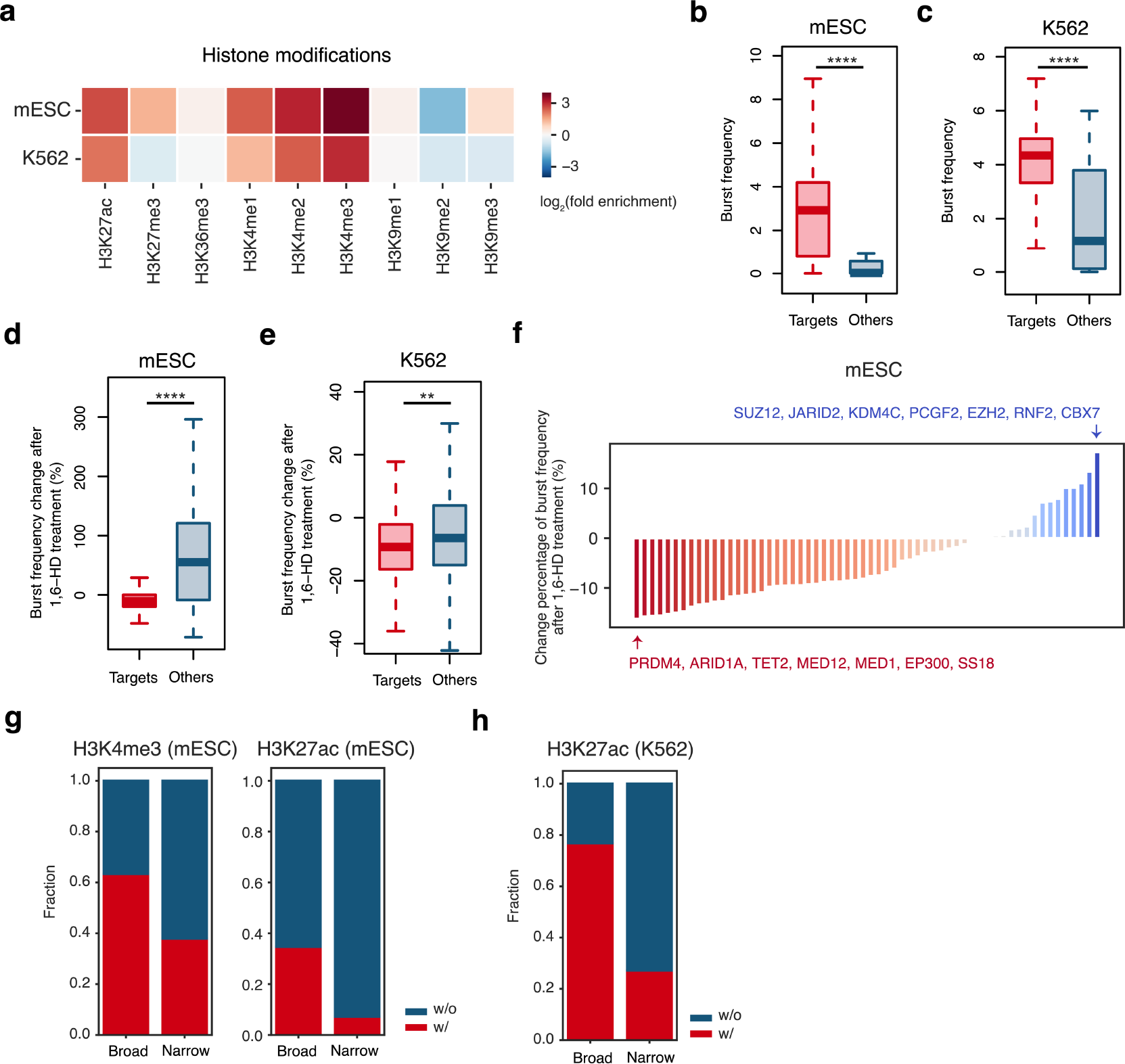
Effect of biomolecular condensates on chromatin activities. **a.** Heatmaps showing histone modification enrichment at CondSig-positive sites in mESC and K562. The colours represent log_2_-transformed fold enrichment of histone modification ChIP-seq signals at CondSig-positive sites relative to the genomic background. Public ChIP-seq data for histone modifications were from Cistrome Data Browser^46^ and filtered as the previous study described^47^. **b, c.** Box plots comparing burst frequency of genes targeted by all CondSig-positive sites and other genes in mESC (**b**) and K562 (**c**). **d**, **e**. Box plots compared the burst frequency change percentage after 1,6-hexanediol treatment of target genes and other genes in mESC (**d**) and K562 (**e**). Significance between groups was evaluated by a two-sided Welch’s *t*-test, ** represents *p*-value < 0.01 and **** represents *p*-value < 1 × 10^−4^. **f.** The bar plots showing change percentages of burst frequency of genes targeted by each individual CondSig. And CondSigs showing the maximum decrease and increase were specially labeled. **g**, **h.** The stacked bar plots showing fractions of broad H3K4me3 or H3K27ac peaks in mESC (**g**) and broad H3K27ac peaks in K562 (**h**) overlapping with CondSig-positive sites.

Notably, several histone modification writers, such as EP300 and KMT2D, were included in the components of identified CondSigs. Given the enrichment of their corresponding histone modifications at CondSig-positive sites (Fig. 5a), we hypothesized that these histone modification writers might exhibit stronger catalyzation activities within biomolecular condensates. We classified each histone modification writer’s ChIP-seq peaks into CondSig-positive and -negative peaks, and observed significantly stronger corresponding histone modification products at CondSig-positive peaks (Supplementary Fig. S6d, e), suggesting the formation of biomolecular condensation can boost the catalyzation activities of histone modification writers. Active modifications, such as H3K4me3 and H3K27ac, typically display narrow peaks (width < 2 kb), while a small proportion also exists as broad peaks (width > 5kb)^27,44^. The establishment of these broad histone modification domains remains unclear, hence we next investigated whether the involvement of their writer in biomolecular condensation could play a role. We transformed two histone modifications’ peaks to domains by merging adjacent peaks not further than 5 kb. Among 1,217 H3K4me3 broad peaks in mESC, 63.3% of them overlapped with KMT2D-associated CondSig-positive sites, while the percentage is only 38.0% for narrow peaks (Fig. 5g). Similar results were observed for the pair of H3K27ac and EP300, not only in mESC, but also in K562 (Fig. 5g, h). These results demonstrated that the involvement of histone modification writers in biomolecular condensates can alter chromatin activity by catalyzing broad histone modification domains.

## Discussion

The field of biomolecular condensate research associated with chromatin has made substantial advancements in recent years. However, identifying the involvement of a CAP in chromatin-associated biomolecular condensate only scratches the surface of its regulatory roles due to the following inherent limitations. Firstly, biomolecular condensates typically comprise multiple components, each potentially contributing different regulatory roles. Secondly, profiling the genomic binding sites of a CAP involved in a biomolecular condensate does not necessarily distinguish its condensation-associated and non-associated genomic loci in a straightforward manner. Therefore, there is an urgent need for specialized experimental methods or bioinformatic tools to provide a detailed genomic landscape of chromatin-associated biomolecular condensates. A recent study introduced DisP-seq^28^, an antibody-independent chemical precipitation assay that maps endogenous DNA-associated disordered proteins at a genomic scale. However, DisP-seq was designed for the broad detection of disordered proteins rather than specifically targeting biomolecular condensates. This could potentially result in both false positives, as not all binding sites of these proteins participate in biomolecular condensates, and false negatives, as disordered protein-guided phase separation is only one mechanism of condensation. Furthermore, DisP-seq cannot identify the exact components present at each locus. In response to these challenges, our study presented CondSigDetector, a computational framework designed to systematically identify CondSigs, *i.e.*, the signatures of condensate-like chromatin-associate protein co-occupancy, and their associated genomic loci. By leveraging the occupancy profiles and condensation-related features of hundreds of CAPs in the same cell type, we can predict the genome-wide loci of biomolecular condensates and the component CAPs of each condensate. Our study both depicted the chromatin properties of the identified CondSigs and experimentally validated the regulatory roles of DDX21, SUPT6H, CTR9 and SUPT5H as components of biomolecular condensates. Our study further delves deeper into the significant effects of chromatin-associated biomolecular condensates on transcriptional bursting and broad active histone modification domains. These findings underscored the critical role that biomolecular condensates play in gene regulation and chromatin activities.

The CondSigs identified in this study provided a comprehensive, global and genome-wide perspective on distinct chromatin-associated biomolecular condensates, paving the way for further exploration of their biological functions and mechanisms. By distinguishing various biomolecular condensates through the unique component CAPs, the CondSigs can not only aid in discovering additional components of known chromatin-associated biomolecular condensates, but also reveal entirely new ones. Furthermore, by pinpointing specific genomic loci targeted by biomolecular condensates composed of CAPs, CondSigs provide valuable insights into how dysregulation of condensation may contribute to disease. This, in turn, could facilitate the design of potential therapeutic strategies. To benefit future research in this area, we have made the CondSigs identified in mESC and K562 publicly available online and provided the source code of CondSigDetector on GitHub to enable the detection in other biological systems.

Despite the significant insights provided by our identified CondSigs, there are some limitations to the predictions. One such limitation is the dependence of CondSig detection on accurate occupancy profiles of CAPs. The absence or poor quality of ChIP-seq data could lead to partial or complete omission of biomolecular condensates. For example, we were able to predict a heterochromatin-related condensate consisting of CBX5, TRIM28 and CBX1 in K562, but not in mESC, due to the unavailability of high-quality ChIP-seq data of these CAPs in mESC. However, with the rapid increase of ChIP-seq data, and the implementation of new techniques for occupancy map capture, we anticipate improvements in the sensitivity of CondSigs detection. Another limitation is the reliance of CondSig detection on specific collaborations among CAPs, which may result in the loss of widespread collaborations in a global context. In this study, we used a threshold of 1.3 for the *z*-score normalized occurrence probability of words in topics to determine the component CAPs of CondSigs. Given the lack of a standard number for components in collaborations, the components listed in CondSig might be incomplete or inaccurate, underscoring the need for further in-depth analysis and experiments to verify the predictions. Finally, a recent study reported that fixation, a common procedure used in X-ChIP, can have diverse effects on biomolecular condensates in living cells^45^. To assess the potential impact of fixation on our prediction results, we selected several component CAPs with additional available data generated by CUT&RUN, a fixation-free technology, to evaluate the concentration levels in CondSigs. We found that, similar to ChIP-seq signals, most component CAPs showed significantly enriched CUT&RUN signals at CondSig-positive peaks (Supplementary Fig. S7), implying that the fixation effect in the X-ChIP procedure is unlikely to significantly impact prediction accuracy. This potential impact could be further mitigated with the rapid accumulation of more CUT&RUN data for CAPs.

## Methods

### ChIP-seq data collection and processing

The ChIP-seq data of CAPs were collected from Cistrome Data Browser^46^ and filtrated using quality control procedures as described in the previous study^47^. In brief, only ChIP-seq data that met at least four out of the five quality control metrics (sequence quality, mapping quality, library complexity, ChIP-enrichment, and signal-to-noise ratio) available in Cistrome Data Browser were kept. In cases where more than one qualified ChIP-seq data were available for a given CAP in the same cell type, all qualified ChIP-seq data were sorted based on quality control metrics, and the top-ranked data was kept.

We downloaded ChIP-seq peak files (in BED format) and signal track files (in bigWig format) from Cistrome Data Brower. Although Cistrome Data Browser stored narrow peaks called by MACS2^48^ for all CAPs, peak window sizes of distinct CAPs could differ significantly. Therefore, to obtain accurate occupancy regions for each CAP, especially CAPs with broad peaks, we first called broad peaks from the signal track using “bdgbroadcall” module of MACS2 (v2.1.3) with default parameters and then merged adjacent peaks within 5 kb. For each CAP, if more than 1,000 newly called peaks were wider than 5 kb, we replaced the original narrow peaks with newly called broad peaks as the accurate occupancy regions.

### Condensation-related annotation for proteins

Human and mouse proteins with reported LLPS capacity were collected from four databases, DrLLPS^6^, LLPSDB^5^, PhaSepDB (two versions, v1 and v2)^3^ and PhaSePro^4^. DrLLPS collected all proteins that could potentially be involved in LLPS, including scaffolds, regulators and clients. However, we only regarded scaffolds as LLPS proteins since DrLLPS contains too many regulators and clients. To create an annotation of LLPS proteins, we merged all LLPS proteins from different sources. Notably, since the number of collected mouse LLPS proteins (61) was much lower than human LLPS proteins (437), we also considered mouse orthologs of human LLPS proteins as mouse LLPS proteins.

Component proteins of MLOs in human and mouse were collected from DrLLPS and PhaSepDB (v1 and v2). Proteins that were assigned to the same MLO in different sources were merged to form a comprehensive list of component proteins for that MLO. Similar to LLPS proteins, mouse orthologs of human proteins assigned to the same MLO was regarded as component proteins of that MLO in mouse.

Pairwise protein-protein interactions were collected from three databases, BioGRID^49^, MINT^50^ and IntAct^51^, only physical associations were kept.

Intrinsically disordered regions of proteins were predicted by MobiDB-lite (v1.0)^52^. This optimized method uses eight different predictors to derive a consensus, which is then filtered for spurious short predictions in a second step. For each protein, if more than 15.3% of its regions were predicted to be disordered by MobiDB-lite, the protein would be regarded as proteins with intrinsically disordered regions. The threshold of 15.3% corresponds to the 20th percentile of disordered region fractions of known human LLPS proteins.

RNA-binding proteins were predicted by TriPepSVM^53^, a method to perform *de novo* prediction based on short amino acid motifs, with parameters “-posW 1.8 -negW 0.2 - thr 0.28”.

### Genome-wide RNA-binding strength

We used genome-wide signals of R-ChIP data, an *in vivo* R-loop profiling approach using catalytically dead RNase H1^54^, to quantify genome-wide RNA-binding strength in K562 cells. Raw sequencing reads from GSE97072^54^ were first aligned to human genome build via default --local mode of Bowtie2 (v2.3.5.1)^55^. Low mapping quality reads (mapping quality < 30) and duplicates were discarded. Then signal tracks were generated using the “genomecov” command in Bedtools software (v2.28.0), and normalized to reads per million mapped reads (RPM).

### Motif scan

Motif scans were performed using FIMO (v5.0.5)^56^ against the JASPAR core 2020 vertebrates database^57^ with the following parameters “--max-stored-scores 1000000”. Motifs with *p*-value ≤ 1 × 10^−5^ were used for the following analysis.

### CondSigDetector workflow

The framework consists of three steps, data processing, co-occupancy signature identification and condensation potential filtration.

In the first step, the framework first splits mouse (mm10) or human (hg38) genome into consecutive 1-kb bins. It then generates an occupancy matrix of CAPs over these 1-kb bins in the given cell type (*n* × *m*), where *n* denotes the number of 1-kb bins and *m* denotes the number of CAPs. The occupancy event of CAP at each genome-wide 1-kb bin is determined by overlapping its ChIP-seq peaks with the given bin. It excludes CAPs with too few occupancy events (those occupying fewer than 500 bins) to eliminate the effect of low-quality ChIP-seq data. And bins with too many occupancy events (occupied by more than 90% of CAPs) are removed to avoid sequencing bias. Additionally, bins in ENCODE Blacklist genomic regions are also discarded.

Identifying co-occupancy signatures from the entire occupancy matrix is a complicated task that can result in the loss of low-frequency signatures in the local context. To address this issue, CondSigDetector first segments the entire occupancy matrix into overlapping sub-matrices iteratively. Each sub-matrix only contains occupancy events of partial highly co-occupied CAPs at partial bins. The segmentation process is as follows: (i) In each iteration, a focus CAP is selected and other CAPs highly co-occupied with the focus CAP are identified. The co-occupancy levels of the focus CAP and the other CAPs are evaluated by using the occupancy events of each other CAP to classify occupancy events of the focus CAP. And then an F_1_ score measuring the accuracy of the classifier is defined as the co-occupancy score and assigned to each other CAP, where a high co-occupancy score implies a high co-occupancy level. In each sub-matrix, only co-occupancy information of the focus CAP and top *q* − 1 other CAPs ranked by the co-occupancy score are kept, where *q* = 50 by default. (ii) After the selection of partially highly co-occupied CAPs, partial bins that are occupied frequently by these CAPs are screened out to further segment the matrix. For *i*-th bin, an occupancy score (*OS_i_*) is defined to evaluate the occupancy level of the given CAPs as:

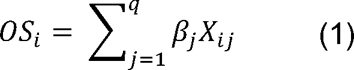

where *X_ij_* ∈ {0, 1} denotes occupancy status of *j*-th CAP at *i*-th bin, and *β_j_* denotes *z*-score normalized co-occupancy score. In each sub-matrix, only *p* bins with *OS_i_* > 0 are kept.

In the second step, each sub-matrix is classified into promoter and non-promoter contexts. Promoters were defined as upstream 3 kb to downstream 3 kb of transcription start sites. CondSigDetector builds a biterm topic model^22^ for each context, treating 1-kb bins as documents and occupied CAPs at those bins as words within documents. By training the model, specific combinations of words can be represented by learned topics, which in turn could be interpreted as co-occupancy signatures representing collaborations of CAPs at chromatin. The biterm topic model is implemented in CondSigDetector using source code from the previous study^22^. As a probabilistic model, the biterm topic model generates two probability distributions, matrix *G_k × q_* representing occurrence probability of *q* CAPs across *k* topics and matrix *G_p × k_* representing occurrence probability of *k* topics across *p* documents.

The topic number, *k*, is a crucial parameter in topic modeling, as it affects the topic distribution. CondSigDetector empirically learns 2∼10 topics for each context and then applies an automatic strategy to select the optimal topic number as described in the previous study^58^. The selection principle was based on the idea that the optimal topic number should distinguish between documents with different topics as much as possible. Hence an optimal topic number should match the following two criteria: (i) The occurrence probability of each topic in different documents should be as different as possible, which is measured by the specificity score (*SS_k_*) calculated for all topics under a certain topic number *k* using Eq. (2). A higher specificity score indicates a better-selected topic number. (ii) The fewer topics that occur in each bin, the better. Such a measurement was defined as a purity score (*PS_k_*) for all topics under a certain topic number *k*, as calculated in Eq. (3). The larger the purity score, the better the selected topic number. Finally, we defined the combination score (*CS_k_*), which is a weighted average of the specificity score and purity score, as calculated in Eq. (4). We selected the optimal topic number from 2∼10 which have the highest combination score.

The specificity score (*SS_k_*) is calculated as

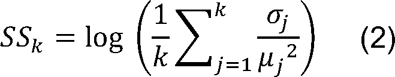

where *σ_j_* and *μ_j_* are the variance and mean, respectively, of the *j*-th column of *G_p × k_*.

The purity score (*PS_k_*) is calculated as

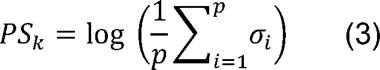

where *σ_i_* is the variance of *i*-th row of *G_p × k_*.

The combination score (*CS_k_*) is calculated as

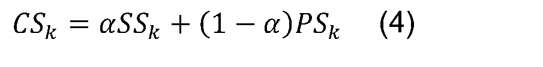

where *α* is calculated as

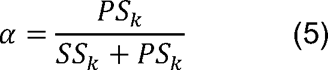

After the selection of optimal topic number *k*, CondSigDetector determined component CAPs of each co-occupancy signature based on matrix *G_k × q_* representing *q* CAPs’ occurrence probability in *k* co-occupancy signatures. In each signature, CAPs with higher *z*-score normalized occurrence probability than a certain threshold (1.3 by default, corresponds to about 90th percentile of the standard normal distribution) were determined as components of the signature, and 1-kb bins occupied by more than 80% of components are defined as signature-positive sites. Co-occupancy signatures with fewer than 3 components and fewer than 200 signature-positive sites are discarded.

In the third step, CondSigDetector screens out CondSigs from all co-occupancy signatures based on the condensation potential of each signature. To evaluate the condensation potential of each signature, we quantify associations between condensation-related features and signature presence at genome-wide bins by performing ROC analysis. Intuitively, the higher condensation-related feature values of occupancy events at signature-positive bins, the higher condensation potential of the signature. In ROC analysis, the positive set is signature-positive bins and the negative set is signature-negative bins. Signature-positive bins were defined in the first step, and signature-negative bins are defined using the following two criteria: (i) The presence of at least *t* CAPs, where *t* = 0.8 × component number of the signature; (ii) The absence of any co-occupancy of components, i.e., count of occupied components of the signature < 2. For each signature, six condensation-related features are calculated according to co-occupancy events of top *q* CAPs (see the first step for the definition of top *q* CAPs): (i) The fraction of occupied CAPs having reported LLPS capacity; (ii) The fraction of occupied CAPs co-occurring in the same MLO; (iii) The fraction of occupied CAPs with predicted IDRs; (iv) The fraction of occupied CAP pairs having protein-protein interactions; (v) The fraction of occupied CAPs predicted as RBPs; (vi) RNA-binding strength of the bin. If at least 3 out of 6 condensation-related features exhibit a positive correlation (AUROC > 0.6) with the presence of the signature (mean AUROC of top 3 features > 0.65), the signature would be identified as CondSigs.

Finally, all CondSigs within the same cell type are pooled and any redundant CondSigs are discarded. Redundancy of CondSigs is measured according to the overlapping level of the top 5 components, these components being ranked by their occurrence probability within the CondSig. We calculate a Jaccard index for all CondSigs using pairwise comparisons, and discard those with a low mean AUROC when the Jaccard index > 0.25. The threshold of 0.25 corresponds to that 2 out of 5 components are identical in the pairwise CondSigs.

### Comparison of BTM and HDP

We built HDP and BTM models on the entire occupancy matrix separately, and compared the quality of learned topics. HDP determines the topic number automatically while BTM asks for a given topic number. So we first built an HDP model and generated *k* topics, then we built a BTM model to generate topics with the given topic number *k.* The quality of each learned topic was evaluated by the coherence score of the top 5 words, a common quality evaluation metric in topic model^22,59^. HDP modeling was implemented by using a Python package “tomotopy”.

### Clustering of component CAPs

We performed a *k*-means clustering for component CAPs in mESC or K562 according to their potentials for self-assembly (PS-Self) or interaction with partners (PS-Part) to undergo phase separation. A recent study employed two machine-learning models, SaPS and PdPS model, to estimate proteins’ potentials and provided SaPS and PdPS ranking scores (ranging from 0 to 1) for the human and mouse proteome. We utilized the SaPS and PdPS ranking scores of component CAPs in mESC or K562 to carry out *k*-means clustering. In the clustering, the number of clusters was set as 4, and the initial cluster centroids were set as (0.8, 0.8), (0.8, 0.4), (0.4, 0.8), (0.4, 0.4), which corresponds to four clusters: “both scaffold and client”, “scaffold-only”, “client-only”, and “none”, respectively.

### Annotation for charged amino acid blocks

We calculated NCPR (net charge per residue) employing a 10-residue sliding window with a step size of 1. This calculation factored in both positively charged amino acids (R, K and H) and negatively charged amino acids (D and E). Windows with NCPR greater than 0.5 or less than −0.5 were defined as charged amino acid blocks, and overlapping blocks were merged.

### Identification of CondSig-positive/negative peaks and domains

To identify CondSig-positive / negative peaks for each component CAP, we classified its ChIP-seq peaks into two groups based on overlapping with positive sites of CondSigs which includes the given CAP as a component. To identify CondSig-positive / negative domains, we transformed its peaks into domains by merging adjacent peaks not further than *n* kb. For component CAPs using narrow peaks as accurate occupancy regions in ChIP-seq data processing procedure, we set *n* = 5, and for component CAPs using broad peaks as accurate occupancy regions, we set *n* = 10. Then domains of each component CAP were classified into CondSig-positive domains and -negative domains based on overlapping with positive sites of CondSigs which includes the given CAP as a component.

### 3D chromatin contact analysis

Public Micro-C data in mESC, ChIA-PET data against SMC1 in mESC, and ChIA-PET data against RNA Pol II in K562 were used in this study. Micro-C contact matrices from 2.6 billion reads were downloaded from GSE130275^30^, and boundary strength for 400-bp resolution calculated by Cooltools^60^ was used for the following analysis. SMC1 ChIA-PET data in mESC were downloaded from GSE57911^33^ and processed with ChIA-PET2^61^. RNA Pol II ChIA-PET loops were directly downloaded from ENCSR880DSH^34^.

### Definition for target genes of CondSig-positive genomic regions

For each genomic region, genes whose promoter overlaps with the given region or has long-range chromatin contacts with the given region were defined as target genes. Long-range chromatin contacts were determined by ChIA-PET data in the corresponding cell type. In this study, SMC1 ChIA-PET data in mESC and RNA Pol II ChIA-PET data in K562 were used.

### Cell culture

Mouse embryonic stem cells (mESC), C57BL/6 strain, were purchased from ATCC (SCRC-1002) and cultured on a feeder layer of mitomycin C (Stemcell, 73272) treated mouse embryonic fibroblast (MEF) in tissue culture flask coated with 0.1% gelatin. The cells were grown in complete mESC medium, which was composed of EmbryoMax DMEM (Millipore, SLM-220-B), 15% (v/v) fetal bovine serum (Hyclone, SH30070.03), 0.1 mM nonessential amino acids (Millipore, TMS-001-C), 1% (v/v) nucleoside (Millipore, ES-008-D), 2 mM L-glutamine (Millipore, TMS-002-C), 0.1 mM β-mercaptoethanol (Millipore, ES-007-E), and 1000 U/mL recombinant LIF (Millipore, ESG1107).

### Cell treatment

1,6-hexanediol (Sigma, 240117) was dissolved in a complete mESC medium at a concentration of 15% (w/v) to make a storage solution. mESC were detached using trypsin, pelleted by centrifuging, and then resuspended in a complete mESC medium. The resuspended cells were transferred into a new gelatin-coated flask and cultured in a 37°C incubator for 1hr to remove the feeder cells. The supernatant cells were collected and washed twice with PBS. After cell resuspending with medium, the 1,6-hexanediol storage buffer was added at a final concentration of 1.5%. The dish was put into the incubator immediately for 30 min, and treated cells were immediately used for CUT&RUN assay.

### CUT&RUN

The CUT&RUN assay was conducted on 0.2 million cells per sample, utilizing the Hyperactive pG-MNase CUT&RUN assay kit (Vazyme, HD102) with slight modifications to the manufacturer’s protocol. Briefly, cells were harvested and incubated for 10 min at room temperature with Concanavalin A-coated magnetic beads, which had been activated prior to use. Following this, the ConA beads bound cells were collected using a magnet and resuspended in 100 µl of antibody buffer containing either 2µl of DDX21 (Proteintech, 10528-1-AP), 4 µl of CTR9 (Novus Biologicals, NB100-1718) or 4 µl of SUPT6 (SUPT6H) (Novus Biologicals, NB100-2582) primary antibody respectively. The samples were then incubated at 4°C overnight on rotator. The next day, cells were washed twice with Dig-wash buffer and resuspended in 100 µl of a premixed pG-MNase Enzyme solution before incubation at 4°C for 1hr with rotation. Following this, the cells were washed twice with Dig-wash buffer and resuspended in 100 µl of premixed CaCl_2_ solution, then incubated for 2hr on ice. Following the stop of the reaction, the cut chromatin was released from cells by incubation at 37 °C for 30 min in the absence of agitation. After centrifuging at 13,400 g for 5 min, the supernatant was collected, and DNA was purified using FastPure gDNA mini columns. The libraries were prepared using NEBNext Ultra II DNA library prep kit (NEB, E7645) with modified amplification condition as 98 °C for 30 sec, 15 cycles of 98 °C for 10 sec and 65°C for 17 sec, and final extension at 65 °C for 2 min and hold at 4°C.

### Single-cell RNA-seq

Single-cell RNA sequencing (scRNA-seq) libraries were prepared using 6,000 mES cells, either in a wild type state or treatment with 1,6-hexanediol at 1.5% for 2 minutes, and K562 cells, either in wild type or treatment with 1,6-hexanediol at 10% for 20 minutes. The libraries were created using the Chromium Single Cell 3’ Library and Gel Bead Kit V3.1 (10x Genomics, Catalog No. PN1000268) to create single-cell gel beads in emulsion (GEM). Following preparation, the libraries were sequenced using the Illumina Novaseq 6000 platform in a 150 bp paired-end mode.

### Immunofluorescence staining

CTR9 antibody and SPT6 (SUPT6H) antibody were labeled with Mix-n-Stain CF488 Antibody labeling kit (Sigma, MX488AS20) and Mix-n-Stain CF568 Antibody labeling kit (Sigma, MX568S20) respectively according to the manufacturer’s instruction. Mouse ES Cells were grown as mentioned above on pre-coated coverslips and fixed with 4% paraformaldehyde solution (Beyotime, P0099) at room temperature for 10 min. permeabilization was performed using 0.5% Triton X-100 (Sigma-Aldrich, 93443) in PBS for 10 min. Cells were blocked with IF blocking solution (Beyotime, P0102) for 1 hr at RT, and subsequently incubated with a 1:100 diluted SUPT5 (SUPT5H) primary antibody (Abcam, ab126592) in QuickBlock dilution buffer (Beyotime, P0262) at 4 °C overnight. Following three washes, cells were incubated with Alexa Fluor 594 goat anti-rabbit secondary antibody (ThermoFisher, A11037) at a concentration of 1: 1000 in PBST for 1 hr at RT. After three additional washes with PBST, cells were labeled with both CF488-conjugated CTR9 and CF568-conjugated SPT6 antibodies at RT for 2hr. After three washes with PBST, the coverslips were mounted onto glass slides using Vectashield medium with DAPI (Vector Laboratories, H-1200) and sealed with nail polish. Images were acquired using a Zeiss LSM 710 confocal microscope with 100 × oil objective and ZEN acquisition software.

### Western blot

Cells were lysed using a lysis buffer (Beyotime, P0013J) supplemented with 1 mM PMSF as a protease inhibitor. Cell lysate was run on a 10% Bis-Tris gel at 70 V for ∼90 min, followed by 120 V until the dye front reached the end of the gel. Proteins were subsequently dry transferred to a nitrocellulose membrane using the iBlot2 western blot transfer system (Thermo Fisher Scientific) under specific conditions: 7 min at 20V for CTR9, and 15 min at 25V for SUPT6H. The membrane was then blocked using 5% non-fat milk in TBST for 1 hr at room temperature with shaking. Primary antibody incubations were performed overnight on a shaker at 4°C with the following dilutions: 1:1,000 of anti-CTR9 (Novus Biologicals, NB100-1718) and 1:1000 of anti-GAPDH (Invitrogen, MA5-15738) in 5% non-fat milk, 1:1,000 of anti-SUPT6 (Novus Biologicals, NB100-2582) and 1:2000 of anti-lamin B1 (Beyotime, AF1408) in Western dilution buffer (Beyotime, P0023). The following day, membranes were washed three times with TBST for 10 minutes each at room temperature with shaking. Secondary antibody incubations were carried out for 2 hours at room temperature using 1:1,000 dilutions of goat anti-rabbit IgG-horseradish peroxidase (HRP) or goat anti-mouse IgG-HRP (Beyotime Technology, A0208, and A0216, respectively) in TBST. Following incubation, membranes were washed three times in TBST for 10 minutes each, and protein bands were visualized by chemiluminescence immunoassay.

### CUT&RUN, single-cell RNA-seq data processing

CUT&RUN reads were first processed using TrimGalore (v0.6.0) to trim adaptor and low-quality reads. Trimmed reads were then aligned to the mouse genome build mm10 or human genome build hg38 using Bowtie2 (v2.3.5.1)^55^ with parameters “--no-mixed --no-discordant --no-unal”. Low mapping quality reads (mapping quality < 30) and duplicates were discarded. Then biological replicates that passed quality control were pooled together. CUT&RUN peaks were called by MACS2 (v2.1.3)^48^. Signal tracks were generated using the “genomecov” command in Bedtools software (v2.28.0), and normalized to reads per million mapped reads (RPM). Single-cell RNA-seq data (10x Genomics) were processed with DrSeq2 (v2.2.0)^62^ and transcriptome-wide transcriptional burst kinetics were inferred using the model from the previous study^42^.

## Data availability

All the CUT&RUN and scRNA-seq data generated in this study have been deposited in Genome Sequence Archive (https://bigd.big.ac.cn/gsa/) under accession CRAXXXXXX and HRAXXXXXX. All predicted CondSigs and associated genomic loci generated in this study are available at https://compbio-zhanglab.org/CondSigDB/index.html.

## Code availability

The computational framework and statistical analysis were made based on shell, Python and R codes. A command-line tool was developed for the implementation of CondSigDetector, main source codes are available at the GitHub repository (https://github.com/TongjiZhanglab/CondSig).

## Acknowledgments

We would like to thank Shuang Hou and Yanhong Xiong for their assistance in statistical analysis and Mengtan Xing for comments on the experimental design. We also thank the staff members of the Integrated Laser Microscopy System at the National Facility for Protein Science in Shanghai (NFPS), Shanghai Advanced Research Institute, Chinese Academy of Sciences, China for sample preparation, data collection and analysis.

## Author Contributions

Y.Z. conceived and designed the research. Z.Y. developed the computational framework and performed computational analysis. Q. W. performed experiments with the help of G.Z and J.Z.. Z.Y, Q. W., and Y. Z. wrote the manuscript.

